# Transient TELC and transmembrane potential in a laser flashed bacteriorhodopsin purple membrane open flat sheet

**DOI:** 10.1101/2024.07.09.602646

**Authors:** James Weifu Lee

## Abstract

The transmembrane-electrostatically localized protons/cations charges (TELC, also known as TELP) model may serve as a unified framework to explain a wide range of bioenergetic phenomenon. Transient TELC and transmembrane potential in a laser flash-energized bacteriorhodopsin (bR) purple membrane (PM) open flat sheet are now better analyzed. Under the Heberle et al. 1994 experimental conditions, the number of bR molecules is now calculated to be 8200 per PM open flat sheet with a diameter of 600 nm. With a single-turnover laser flash intensity of 3 mJ/cm^2^ to photoexcite 10% of the bR molecules, the number of laser flash-induced peak TELC density is calculated to be 2900 per µm^2^ of PM, which translates to a peak transient transmembrane potential of 50 mV. The bR protonic outlet protrudes into the liquid phase outside the putative “potential well/barrier”. The observation is in line with the TELP model; but does not support the “potential well/barrier” model. The author encourages research on more relevant protonic cell systems that have transmembrane potential with TELC comprising excess positive charges at one side and excess anions at the other side of the membrane.

## Introduction

According to the transmembrane-electrostatically localized protons/cations charges (TELC) model [1] [2] [3], transmembrane potential (Δ*ψ*) is a function of TELC surface density, as shown in the following protonic membrane capacitor-based equation with a voltage unit (V in volts):

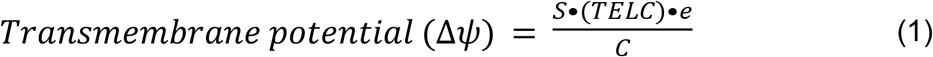

where *C/S* is the specific membrane capacitance per unit surface area; *TELC* is the number of positive charges per unit membrane surface area, which is the sum of transmembrane-electrostatically localized protons (TELP) and transmembrane-electrostatically localized non-proton cations after cation exchange with TELP; and *e* is a proton (or cation) charge of 1.60 x 10^−19^ Coulomb.

Accordingly [1] [2] [3], when a transmembrane proton pumping process which is an electrogenic (i.e., electrically non-compensated) charge transport across a bacteriorhodopsin purple membrane open flat sheet is energized by a single-turnover pulsed laser flash [4] [5], a transient transmembrane potential (Δ*ψ*) will form and then decay as the pumped excess charges returning to the other sides of the membrane. The formation of a transient transmembrane potential (Δ*ψ*) is resulted from a protonic capacitor formation with transmembrane-electrostatically localized protons (TELP) on the positive (extracellular) side and with transmembrane-electrostatically localized hydroxide anions on the negative (cytoplasmic) side along a bacteriorhodopsin purple membrane open flat sheet, at least, transiently. That is, during its protonic capacitor formation, according to Eq. 1, there is at least a transient transmembrane potential (Δ*ψ* ≠ 0), in contrast to Silverstein’s claim of no “transient non-zero Δψ”: “where Δψ = 0” [6] [7].

In this article, we will calculate the bacteriorhodopsin (bR) population density in a bacteriorhodopsin purple membrane open flat sheet that was tested in the experiment of Heberle et al. 1994 [5]. Then, the transient TELC population and transmembrane potential in a laser flashed bacteriorhodopsin purple membrane open flat sheet will be calculated. The analyzed results will be discussed with commentary in the context of Silverstein’s recent argument [7] and the putative “potential well/barrier” model proposed by Junge and Mulkidjanian [8, 9] and advocated by Silverstein [7].

### Calculation of transient TELC and transmembrane potential in a laser flashed bacteriorhodopsin purple membrane open flat sheet

The experiment of Heberle et al. 1994 [5] employed a piece of well-characterized purple membrane (PM) of *H. salinarium*, which is “a two-dimensional crystalline array of the integral membrane protein bacteriorhodopsin with lipids” that is also called as a bacteriorhodopsin purple membrane open flat sheet. The natural PM composed of 75% bR (w/w) and 25% lipid (w/w) [10]. Each bacteriorhodopsin (bR) molecule consists of approximately seven helical segments transversing the membrane [11]. The crystal structure of PM which naturally forms is a trimer of bR molecules within one hexagonal unit cell (a = 62 Å) as shown in the AFM images (Figure 1) [12]. That is, the bR trimers (encircled with the triangles) are arranged in a hexagonal lattice with unit cell dimension of ∼62 Å [12]. Using this information, as shown in Table 1, we have now calculated the number of bR molecules to be 8200 bR molecules per PM open flat sheet with a diameter of 600 nm that was tested in the experiment of Heberle et al. 1994 [5].

**Table 1.** Calculation for the number of bacteriorhodopsin (bR) molecules per purple membrane (PM) disk (with a diameter of 600 nm) employed in Heberle et al. 1994 experiment. The laser wavelength was 532 nm, and its pulse width was 8 ns. A pulsed laser flash (8 ns) energy density of 3 mJ/cm^2^ was used to photoexcite only about 10% of the bR molecules to ensure single-turnover conditions [4]. Peak transient transmembrane potential (Δ*ψ*) was calculated from peak TELC density through Eq. 1 using specific membrane capacitance C/S of 9 mf /m^2^ based on experimentally measured cell membrane capacitance data [13].

| Laser flash intensity | bR molecules per PM disk | Peak TELC per PM disk | Peak TELC per $\mu\text{m}^2$ of PM | Peak transient transmembrane potential ($\Delta\psi$ in mV) | Lifetime ($\mu\text{s}$) of transient TELC and transmembrane potential |
| --- | --- | --- | --- | --- | --- |
| 0 | 8200 | 0 | 0 | 0 | NA |
| 3 $\text{mJ}/\text{cm}^2$ | 8200 | 820 | 2900 | 50 | 152 |
| 6 $\text{mJ}/\text{cm}^2$ | 8200 | 1640 | 5800 | 100 | 152 |

**Figure 1.**
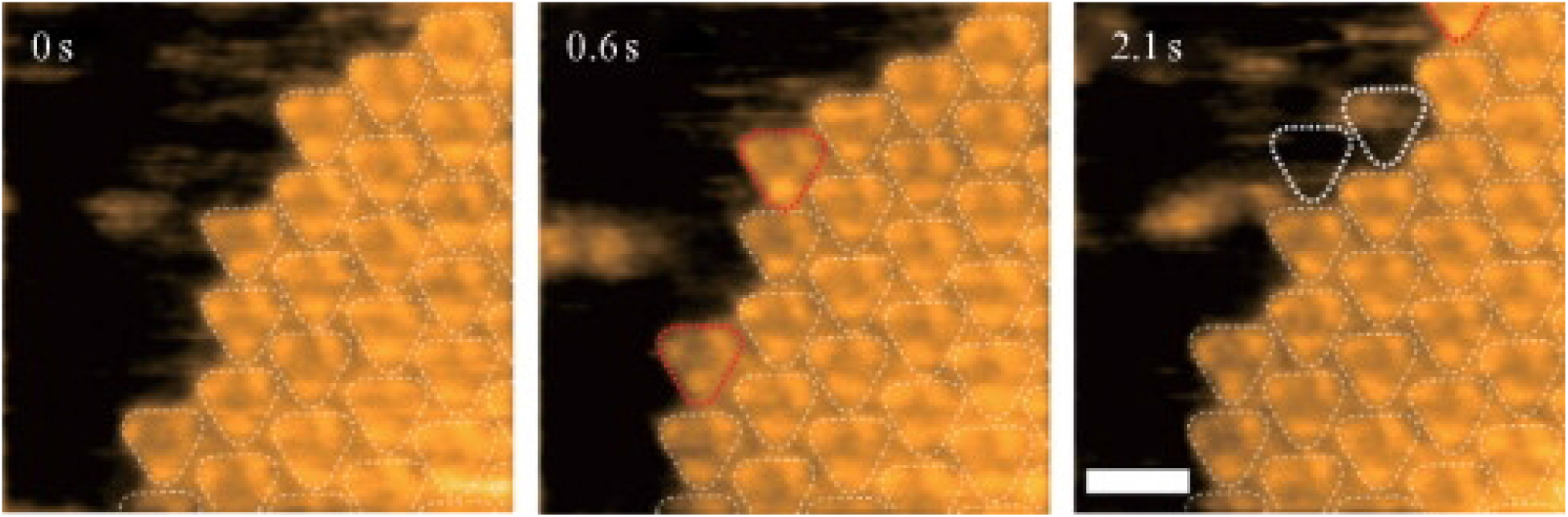
AFM image of the crystal edge of bacteriorhodopsin (bR) in purple membrane. The bR molecules encircled by the dotted lines golden triangles indicate the bound bR trimers. The AFM images were taken at 3.3 frames/s (scale bar, 10 nm). The bR molecules encircled by the red dotted lines (at 0.6s) indicate newly bound bR trimer. The white triangles (at 2.1 s) indicate the previously bound trimers. The bR trimers (encircled with the triangles) are arranged in a hexagonal lattice with unit cell dimension of ∼62 Å. Reproduced with permission from (Yamashita et al., 2009) [12].

In the Heberle et al. 1994 experiment, the bR PM open flat sheet was transiently energized by use of a single-turnover pulsed laser flash. The laser wavelength was 532 nm and pulse width was 8 ns. A pulsed laser flash (8 ns) energy density of 3 mJ/cm^2^ was used to photoexcite only about 10% of the bR molecules to ensure single-turnover conditions [4]. Therefore, upon a single-turnover laser flash energization at the intensity of 3 mJ/cm^2^, it will transiently generate 820 charge-separated pairs of bRH^+^ and bROH^*−*^ across the bacteriorhodopsin PM disk (diameter 600 nm), which constitutes 820 TELC per PM disk that is equivalent to a peak TELC density of 2900 per µm^2^ of PM. From the peak TELC density of 2900 per µm^2^ of PM, we employed Eq. 1 using specific membrane capacitance *C/S* of 9 mf /m^2^ based on experimentally measured cell membrane capacitance data [11] and calculated the peak transient transmembrane potential to be approximately about 50 mV (Table 1).

If the laser flash intensity is doubled to 6 mJ/cm^2^, then the peak TELC density may double to 1640 TELC per PM disk (which is equivalent to a peak TELC density of 5800 per µm^2^) that will give rise to a peak transient transmembrane potential of about 100 mV as shown in Table 1.

As listed in Table 1, the lifetime of the transient transmembrane potential is expected to be the same as the associated transient TELC lifetime (152 µs) that was calculated from the experimental data of Heberle et al. 1994.

### Transient TELC activity along a laser flash-energized bacteriorhodopsin purple membrane open flat sheet

Therefore, as previously discussed [3], the bR PM open flat sheet in liquid water can be regarded as a special protonic capacitor disk with its edge (rim) connected through a protonic conductor (water) as shown in Figure 2A. Thus, when protons are actively pumped across the membrane through the membrane-embedded bR photochemistry mechanism from the cytoplasmic side to the extracellular side, it will create excess protons (bRH^+^) on extracellular surface while leaving equal number of excess hydroxide anions (bROH^−^) on the cytoplasmic surface.

**Figure 2.**
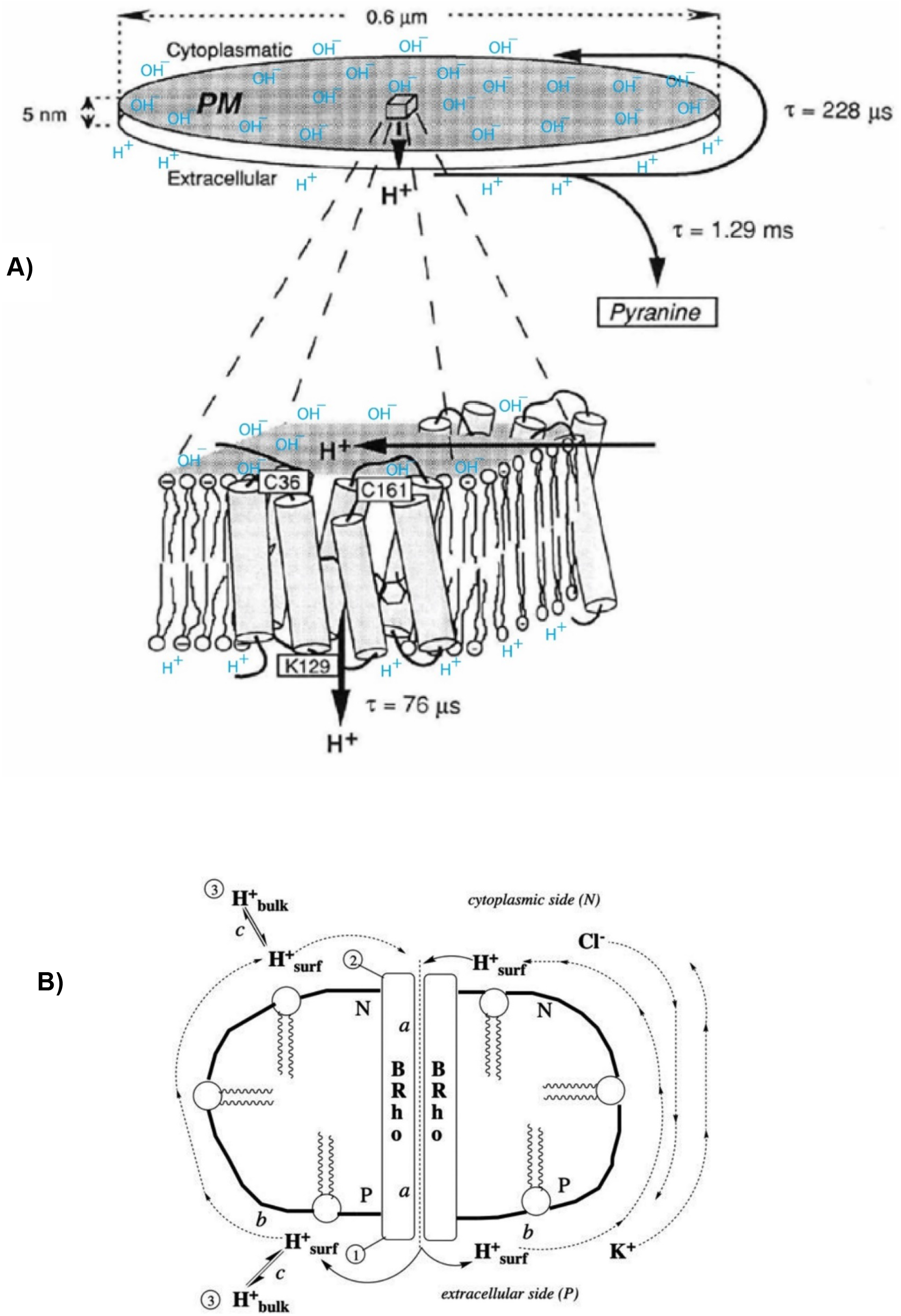
**A)** Formation of a transient protonic capacitor with transmembrane-electrostatically localized protons (blue, H^+^) at the liquid-membrane interface on the extracellular side (bottom) and the localized hydroxide anions (blue, OH^−^) on the cytoplasmic side (top) along a bacteriorhodopsin purple membrane open flat sheet. Adapted from Ref [3], which is modified from Ref [5]. **B)** Silverstein’s particle-like drawing with a single bR molecule that he apparently intended to represent a purple membrane open flat sheet (that is now known to have 8200 bR molecules): (a)light-driven pumping across the membrane, through bacteriorhodopsin “BRho”; (b) surface diffusion; and (c) release from the surface into the bulk phase. The three proton detection sites are (1) bR lys129 at the *P* surface; (2) bR cys36 and cys161 at the *N* surface; and (3) aqueous pyranine. Reproduced with permission from Silverstein 2023 [7].

The charge-separated pairs of excess protons (bRH^+^) and excess hydroxide anions (bROH^−^) created by the pulsed laser energized bR photophysical chemistry across the PM represent the initial (peak) population of the laser pulse-induced TELP (TELC), which can translate to a peak transient transmembrane potential through Eq. 1 as shown in Table 1.

Therefore, according to the protonic capacitor model [1] [2] [3], the excess protons in this case will be held, at least transiently, at the liquid-membrane interface along the extracellular surface by the transmembrane-electrostatic attractive force from the excess hydroxide anions at the liquid-membrane interface along the cytoplasmic surface as illustrated in Figure 2A. Subsequently, the excess protons will rapidly move along the liquid-membrane interface on the extracellular surface and through the PM rim to return to the cytoplasmic side; meanwhile the excess hydroxide anions can also quickly translocate on the cytoplasmic surface towards the rim of the bR PM flat sheet. Also notice that, from the negative charge point of view, hydroxide anions (protonic holes) are transferred in the opposite direction of proton conduction. Consequently, the translocation of the excess hydroxide anions on the cytoplasmic surface towards the rim of the bR PM open flat sheet is equivalent to the translocation of excess protons from the rim towards the center of the cytoplasmic surface in filling the “protonic holes” there. When the excess protons finally recombine with the excess hydroxides in forming water molecules, the transient transmembrane potential (Δψ) decays to zero.

Therefore, to the observers who monitor the translocated protons using some localized protonic sensors (such as pH-indicating dye fluorescein) along two sides of the bR PM open flat sheet and using certain delocalized pH probe (pyranine) in the bulk liquid phase, they would observe that the excess protons pumped through bR appear moving along the extracellular membrane surface and then passing through the liquid around the rim to the cytoplasmic surface, but without getting into the bulk liquid phase as shown in Figure 2A. These predicted features were exactly shown up in the bR PM open flat sheet experiment that was elegantly performed by Heberle et al. [5] where they employed a pulsed laser flash to suddenly energize bacteriorhodopsin-driven H^+^ pumping and monitored the flash-induced transient absorption changes of the protonic molecular probes. Therefore, as recently reported [3], the prediction from the TELP model can now well explain the experimental results of Heberle et al. [5] that suggest “protons can efficiently diffuse along the membrane surface between a source and a sink (for example H^+^-ATP synthase) without dissipation losses into the aqueous bulk”.

### Discussion and commentary in responding to Silverstein 2023 argument

Recently, Silverstein 2023 [7] tried again to defend his notion of “where Δψ = 0” (9)using a bR membrane particle-like drawing (less than 10 nm in size as judged by the size of liquid molecules with a single bR molecule?) as shown in Figure 2B to represent the 600 nm diameter flat PM disk (Figure 2A) of Heberle et al. 1994 [5]. Note, the 600 nm diameter (geometric size) of a flat PM disk and the number of 8200 bR molecules per disk (see Table 1) are relevant parameters that may affect the TELC-associated transport activities and the lifetime and amplitude of transient transmembrane potential. Probably trying to favor his argument, Silverstein’s drawing (Fig. 2B) shows only a single bacteriorhodopsin molecule with a few lipid molecules. According to the length of a lipid molecule which is known to be about 2 nm (and a typical biomembrane thickness of 4 nm), his drawing (Fig. 2B) seems to have the appearance of inappropriately portraying the diameter 600-nm PM disk (containing 8200 bacteriorhodopsin molecules) as a single bR membrane particle with a size of 10 nm?. Since his article [7] does not mention anything about the 600 nm diameter of the flat purple membrane disk, his 10-nm particle-like drawing with a single bR molecule (Fig. 2B) could give misimpression about the Heberle et al 1994 experiment [5] and cause confusion in the field.

Silverstein’s recent argument [7] centers on “the circuit is ‘shorted out’ by the free flow of ions between the two sides of the membrane, around the rims of the membrane fragments.” However, Silverstein’s own statement “Once a proton is released on the P side and a transient Δψ is generated, ions in solution will respond: K^+^ will move from the P side toward the N side, and Cl^−^ will move in the opposite direction” professes “a transient Δψ is generated”.

There is no evidence for his proposed vectorial ion flow: “K^+^ will move from the P side toward the N side, and Cl^−^ will move in the opposite direction” at the PM flat sheet rim as he tried to portray with his 10-nm particle-like drawing with a single bR (Figure 2B) [7]. Even if his proposed vectorial ions flow were true, that would still have to be driven by an electrical field from the transient transmembrane potential Δψ, in accordance with the Fick’s laws of diffusion; it again indicates the presence of transient membrane potential (transient peak Δψ = 50 mV as calculated in Table 1) in contrast to his notion [6] of “where Δψ = 0”.

Silverstein [7] repeatedly argued that the photo-driven bR proton pump “is 56,000 times slower than proton diffusion in bulk aqueous solution”. This argument is irrelevant, since the pulsed laser-induced formation of transient membrane potential (transient peak Δψ = 50 mV) is the first (primary) step (1) because of the photo-driven bR proton pump, which is an active transport process. All the others are the passive processes, including (2)proton translocation along the liquid-membrane interface (and around the PM flat sheet rim) and (3) slow (1.29 ms) partial (17%) proton releasing into the bulk liquid phase likely as a result of cation-proton exchange associated with ion and proton diffusion in bulk aqueous solution [3]. They (2 and 3) <u>happen later</u> than the primary active transport step (1) since the passive transport processes (2 and 3) must be driven, at least transiently, by the pulsed laser-induced bR photophysical proton-pumping charge separation process (1) that creates a transient membrane potential (transient Δψ ≠ 0) as the driving force.

By now, readers can probably identify the mistakes in Silverstein’s arguments “Given *D* = 2 nm^2^/ns for K^+^ and Cl^−^ in bulk water [14], it would take only 0.002 μs (= (5 nm)^2^/(6 × 2 nm^2^/ns) = 2 ns) for one of these ions to cross the 5 nm around the rim of the membrane fragment from the P and N side … So, light-driven H^+^ pumping via bacteriorhodopsin across the purple membrane fragment is not electrogenic; in the 76 μs it takes to pump a proton across the membrane, it is easily electrically compensated by K^+^ and Cl^−^ diffusion around the fragment rim. Thus, one would not expect to find even a transient non-zero Δ*ψ* in this system”.

Note, the pulsed laser-induced 76-μs photo-driven bacteriorhodopsin active proton pump and transient transmembrane potential formation is the <u>primary (active) transport</u> step (1) that drives the sequence of the subsequent transport events (2 and 3). The <u>passive transport</u> processes, including (2) proton translocation along the liquid-membrane interface and around the rim plus (3) slow (1.29 ms) partial (17%) proton releasing likely through cation-proton exchange associated with ions and proton diffusion in bulk aqueous solution [3], all happen later after the laser pulse-induced primary (active) transport step (1). In his arguments above, Silverstein seems to assume that the passive transports (2 and 3) could occur simultaneously or before the laser pulse-induced primary (active) transport step (1), which seems to be his fundamental mistake. Therefore, readers may now be able to see four mistakes in Silverstein’s arguments [7]:

1. Improperly mixed up the sequence of events from the primary event of the 76-μs photo-driven bR <u>active</u> proton pump and transient transmembrane potential formation (1) to the <u>passive</u> transport processes including: (2) proton translocation along the liquid-membrane interface and around the rim plus (3) slow (1.29 ms) partial (17%) proton releasing associated with ions and proton diffusion in bulk aqueous solution [3];
2. Wrongly assumed a passive vectorial “flow of K^+^ and Cl^−^ cross the 5 nm around the rim” to occur simultaneously or before the laser pulse-induced 76-μs bR proton-pump event that resulted in formation of a transient membrane potential (peak transient Δ*ψ* = 50 mV as calculated in Table 1);
3. Inaccurately assumed a passive vectorial “flow of K^+^ and Cl^−^ cross the 5 nm around the rim” could <u>instantaneously (𝒯=0)</u> reduce transmembrane potential to zero;
4. Wrongly claimed his “2 ns” for a passive vectorial ion flow to cross the 5 nm around the rim as a decay lifetime for the transient transmembrane potential; even if the claimed “2 ns” decay time were true, it would still admit the transient transmembrane potential (transient Δ*ψ* ≠ 0) during its lifetime.

Note, the decay lifetime (𝒯) for a transient transmembrane potential (transient peak Δψ = 50 mV) is expected to be correlated with the amount of time that the excess protons at the extracellular side take to translocate from the center area of the 600-nm diameter bR PM open flat sheet through the rim to the center area of the cytoplasmic side to recombine with the excess hydroxides to form water molecules. According to the Heberle et al. 1994 experimental data [5] assuming a mean distance for the two-dimensional diffusion of 240 nm (Fig. 5), we estimated the decay lifetime (𝒯) for the transient membrane potential (transient Δψ ≠ 0) to be about 152 μs (𝒯 = 228 μs – 76 μs) as listed in Table 1.

Silverstein [7] recently also tried to argue for the presence of “the porins and ion channels” in the 600-nm diameter bR PM open flat sheet (Fig. 5) employed in the Heberle et al. [5] experiment. That argument is also moot since the PM open flat sheet is composed of 75% bR (w/w) and 25% lipid (w/w) [10]; there is no evidence for his claimed “porins and ion channels” in the 600-nm diameter bR PM open flat sheet that was used in the Heberle et al. 1994 experiment [5].

As recently reported [3], it can be shown through the ohms law: the voltage *V* = *I*·*R* ≠ 0 where *I* is the total current of protonic and ionic conduction through the bR PM rim plus through “the porins and ion channels” (if present); and *R* is the protonic and ionic resistance of the liquid medium which has a non-zero resistance. Therefore, his argument [7] and notion [6] of “where Δψ = 0” for a laser-energized flat bacteriorhodopsin membrane sheet is again just a misunderstanding. The argument shall now be over; Under the experimental conditions of Heberle et al. 1994 experiment [5], the peak transient transmembrane potential (Δψ) has now been numerically calculated to be 50 mV (Table 1) based on the numbers of the charge-separated pairs of excess protons (bRH^+^) and excess hydroxide anions (bROH^−^) created by the laser-energized bR photophysical chemistry across the PM sheet.

### Protonic outlet of bR structure rejecting the “potential well/barrier” model

Silverstein [7] presented a quite interesting plot for the putative “potential well/barrier” model’s Gibbs free energy profile energy for moving protons from the water/decane interface to the bulk aqueous phase (Figure 3A). He wrote “Δ*G* for the depth of the potential well near the water/decane interface (4–8 *RT* at 1–2 Å) was calculated by ab initio free energy molecular dynamics [15]. The activation barrier, Δ*G*^°‡^ ≈ 20–30 *RT* is calculated from Arrhenius/Eyring plots [15] [16].”

**Figure 3.**
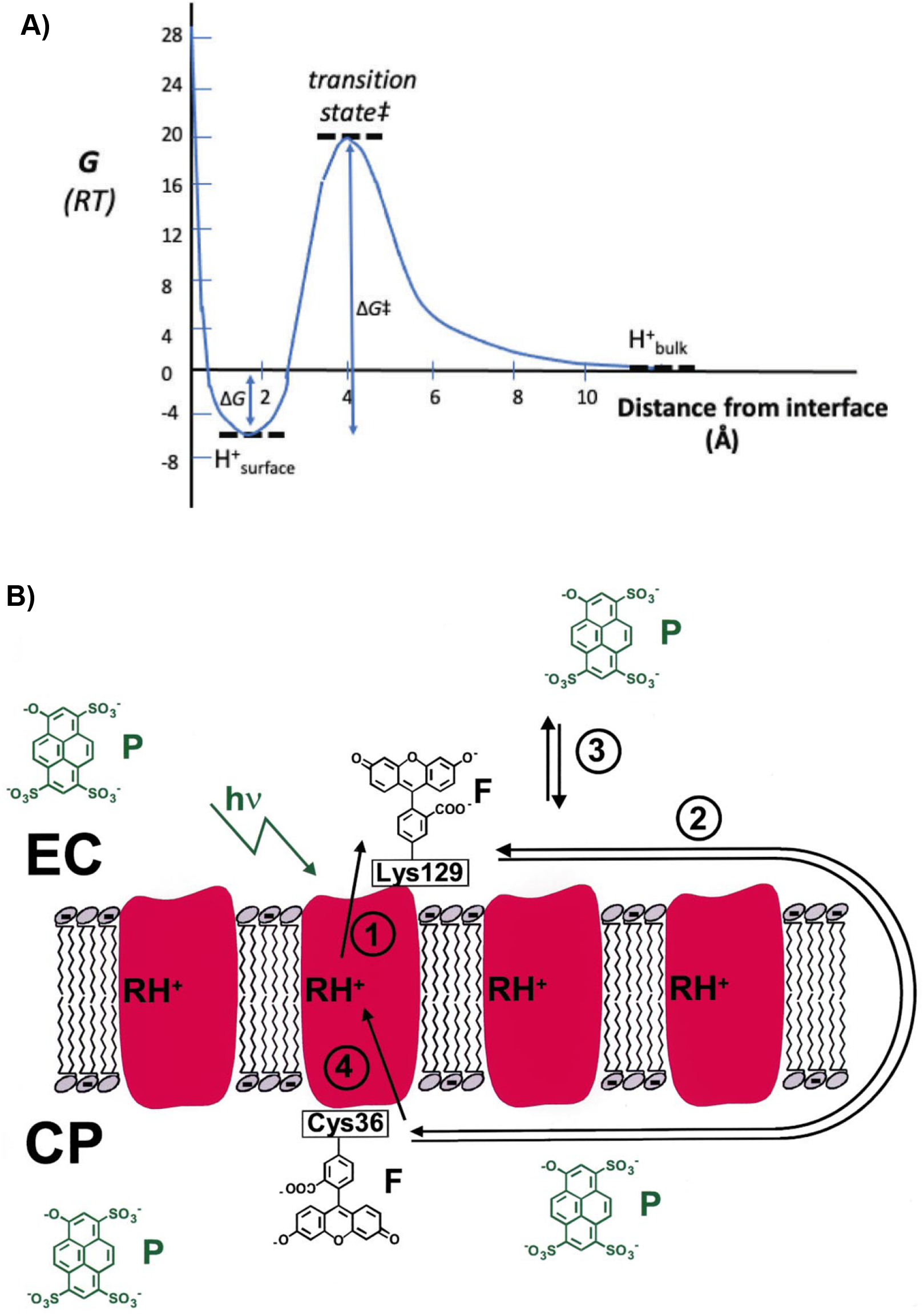
**A)** Free energy profile energy for moving protons from the water/decane interface to the bulk aqueous phase. Δ*G* for the depth of the potential well near the interface (4–8 *RT* at 1–2 A) was calculated by ab initio free energy molecular dynamics [15]. The activation barrier, Δ*G*^°‡^ ≈ 20–30 *RT* is calculated from Arrhenius/Eyring plots [15] [16]. Reproduced with permission from Silverstein 2023 [7]. **B)** Proton transfer reactions across bR and along the PM surface. Sketch of bR molecules (RH^+^, protonated Schiff base) in the PM with fluorescein (F) covalently bound to the amino acid K129 at the extracellular side and to C36 at the cytoplasmic side. Pyranine (P) resides in the aqueous bulk phase. Reproduced with permission from Ref. [10].

At the experimental temperature of 20 ^o^C (293 K) [5], Silverstein’s claimed “Δ*G*^°‡^ ≈ 20–30 *RT* “ (+ 25 *RT* ?) in the liquid water phase at the location of 0.4 nm away from the decane surface would be equivalent to an activation barrier of about 60.9 kJ/mol; that is likely to be flawed or questionable since he has never explained how could it be possible for water molecules to form such a high activation barrier that is far much higher than their hydrogen bond energy? According to an independent study [17], the water hydrogen bond Gibbs energy *ΔG*^*0*^ is 2.7 kJ/mol (with *ΔH*^*0*^ = 7.9 kJ/mol and *TΔS*^*0*^ = 5.2 kJ/mol). For the mechanism of proton mobility, the activation energy *E*_*A*_ is known to be 11.3 kJ/mol [17] [18], in contrast to Silverstein’s claimed “Δ*G*^°‡^ ≈ 20–30 *RT* “ (around 60.9 kJ/mol).

Furthermore, even if assuming the putative “potential well/barrier” model would be true, it still could not explain the relevant bioenergetics. As shown in Figure 3A, the bottom of its putative Gibbs free energy potential well (−6 *RT*, if true) would be located about 0.15 nm away from the decane surface; and the peak of the activation barrier (+25 *RT*, if true) would be located at 0.4 nm away from the decane surface. Consequently, if the “potential well/barrier” model (Figure 3A) were correct, it would imply that the protonic outlet of bR molecular structure would have been located precisely inside the “potential well” that would be within 0.4 nm from the alkane surface of the membrane. Otherwise, the bR protonic outlet (which protrudes into the liquid phase at least about 1 nm away from the hydrophobic core membrane surface) will be outside the putative “potential well/barrier”. If the protonic pump outlet protrude more than 0.6 nm away from the hydrophobic core membrane surface (so that it is outside the putative potential well/barrier), the “potential well/barrier” model (even if exists) would not work.

The proton (H^+^) transfer processes from the active center of bR to the extracellular side (Fig. 3A, reaction 1) and from there along the membrane-water interphase to the cytoplasmic side of bR (reaction 2) are kinetically and spatially resolved [10]. The pH indicator fluorescein (F) that was bound to K129 at the extracellular surface of bR demonstrated the bR protonic outlet activity at the position well above the lipid head groups, which is at least 1 nm away from the alkane surface of the membrane. As shown in Figure 3B, bR protonic outlet (K129) apparently protrudes into the liquid phase at least about 1 nm away from the lipid bilayer’s alkane membrane surface. That is, the bR protonic outlet is located at the liquid phase outside the “potential well/barrier” (Figure 3A).

This observation (Figure 3), which is consistent with the known bR structure and function [19] [11] [20], is well in line with the TELP model; but it does not support the putative “potential well/barrier” model. As more clearly shown in the X-ray crystallographic structure of bacteriorhodopsin (Figure 4A), its protonic outlet protrudes and injects proton (H^+^) into the bulk liquid phase of the extracellular space at least about 1 nm away from the membrane surface [20]. Another independent study [11] (Figure 4B) also showed that the membrane-embedded bacteriorhodopsins translocate proton (H^+^) into the bulk liquid phase of the extracellular space at least about 1 nm away from the membrane surface. Note, the Heberle et al. 1994 experiment [5] demonstrated the laser-induced bR proton release (𝒯 = 76 µs) at its K129 which apparently protrudes into the bulk liquid phase (bottom left, Figure 4B). Therefore, the structure and function of bacteriorhodopsin (Figure 4) quite clearly rejects the putative “potential well/barrier” model (Figure 3A).

**Figure 4.**
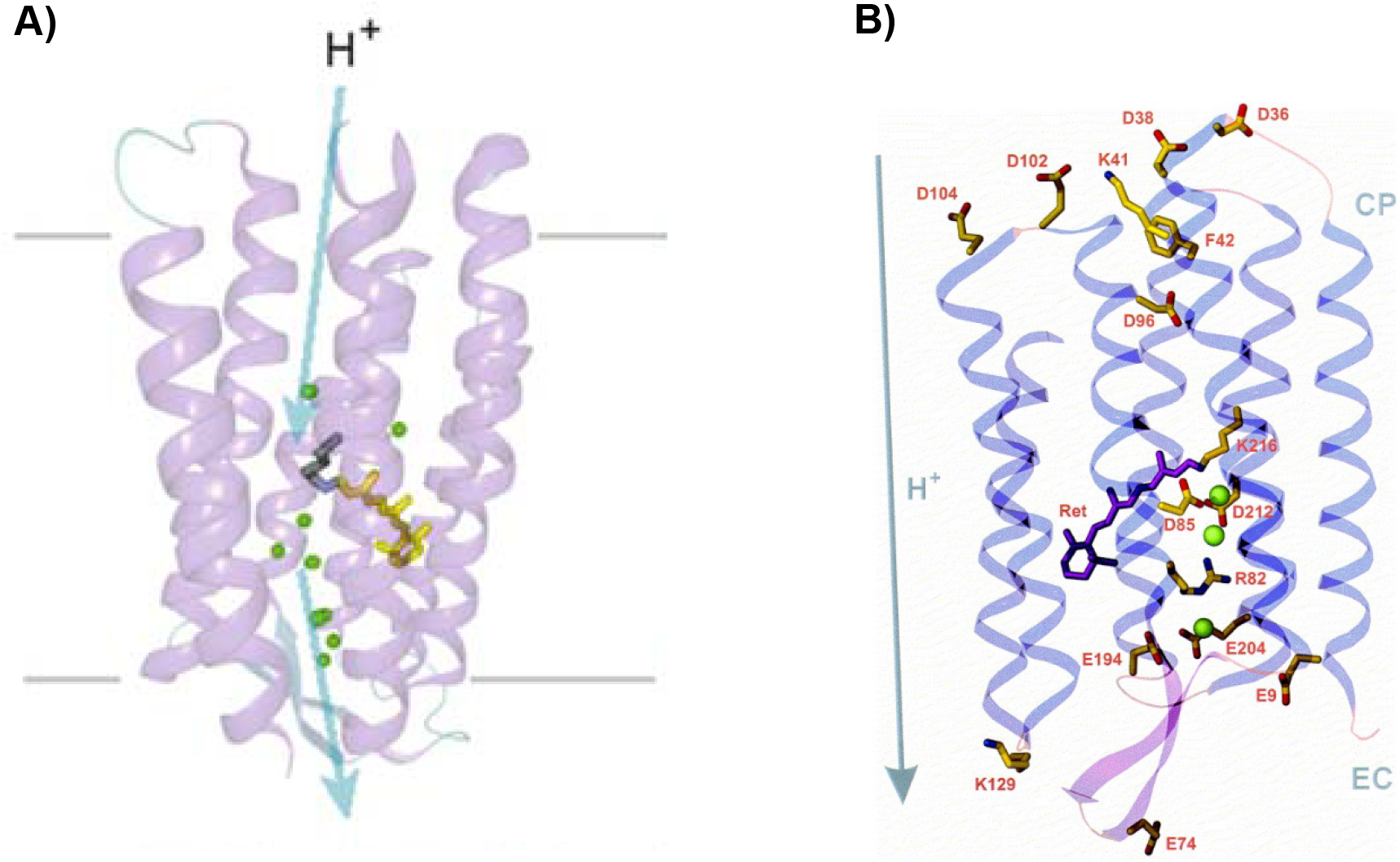
**A)** X-ray crystallographic structure of bacteriorhodopsin shows its protonic outlet injecting proton (H^+^) into the bulk liquid phase of the extracellular space at least about 1 nm away from the membrane surface. Reproduced with permission from Ref. [20]. **B)** Structural model of bacteriorhodopsin based on a resolution of 2.3 Å ([19]; data retrieved from the protein data bank, PDB entry: 1BRX). View is approximately parallel to the membrane plane. The thickness of the surrounding membrane is about 40 Å. The protein backbone is shown as ribbons. The chromophore retinal (Ret) and amino acids are represented as sticks. Three water molecules are shown as spheres. The arrow indicates the direction of proton translocation. Reproduced with permission from Ref. [11]

In addition to the structure of bR protonic outlet of bR that apparently rejects the “potential well/barrier” model, many other known protonic pump outlets such as those of complexes I, III and IV in mitochondria also apparently protrude into the liquid phase at least 1-3 nm away from the membrane surface [21] [22] [23]; and they thus do not support the “potential well/barrier” model either. Therefore, it is now again quite clear that the putative “potential well/barrier” (Figure 3A) as proposed by Junge and Mulkidjanian [8, 9] and advocated by Silverstein [7] does not really exist, or the putative potential well/barrier (even if exist) is irrelevant to explaining the results of Heberle et al. 1994 experiment [5].

Therefore, this author encourages more scientific research efforts on more relevant protonic cell systems that have transmembrane potential with TELC comprising excess positive charges at one side and excess anions at the other side of the membrane.

## Acknowledgement

The author thanks Dr. Joachim Heberle and Dr. Norbert Dencher for providing more detailed information about their 1994 experimental conditions including the information about the bacteriorhodopsin molecular population density in the bacteriorhodopsin purple membrane open flat sheet and their pulsed laser property and operating conditions that made some of the analyses reported in this article possible. The author thanks Todd Silverstein for the stimulating discussions through numerous emails. The author also thanks the anonymous peer reviewers for their highly valuable and constructive review comments that made this article better.

## AUTHOR INFORMATION

## Author contributions

Lee designed and performed research, analyzed data, and authored the paper.

## Data availability

All data generated or analyzed during this study are included in this article and in the cited references.

